# Failure of VITEK2 to reliably detect *vanB*- mediated vancomycin resistance in *Enterococcus faecium*

**DOI:** 10.1101/2020.11.11.379180

**Authors:** Sarah V. Walker, Martina Wolke, Georg Plum, Robert E. Weber, Guido Werner, Axel Hamprecht

## Abstract

**Objectives:** The increasing prevalence of vancomycin resistant enterococci (VRE) necessitates a reliable detection of VRE especially for low level resistance mediated by *vanB* in *Enterococcus faecium*. In this prospective study we analyzed if *vanB* mediated vancomycin resistance can be reliably detected by Vitek2.

**Methods:** 1344 enterococcal isolates from routine clinical specimens were tested by Vitek2 (bioMérieux, Nürtingen, Germany). Additionally, a bacterial suspension (0.5 McFarland) was inoculated on a chromID VRE screening agar (bioMérieux) and incubated for 48 hours. If vancomycin was tested susceptible by Vitek2 but growth was detected on the screening agar a PCR for *vanA/vanB* was performed (GeneXpert *vanA/B* test kit, Cepheid, Frankfurt, Germany). MICs of vancomycin susceptible by Vitek but *vanA/B* positive isolates were determined before and after cultivation in a broth with increasing concentration of vancomycin.

**Results:** 156/492 of *E. faecium* were VRE, predominantly *vanB* (87.0%) of which 14 were not identified as VRE by Vitek2 (sensitivity 91.0%). The majority (9/14) demonstrated high-level MICs by broth dilution. Even after exposure to increasing vancomycin concentrations MICs remained nearly identical. Three of the undetected isolates demonstrated initial growth on chromID VRE, after the vancomycin exposure additional 7 isolates demonstrated growth on chromID VRE.

**Conclusions:** Vitek2 fails to detect *vanB* mediated vancomycin resistance consistently, especially but not limited to low-level resistance. As this may lead to treatment failure and further dissemination of vanB VRE, additional methods (e.g. culture on VRE screening agar or PCR) are necessary to reliably identify *vanB*-positive enterococci in clinical routine.

## Introduction

Over the last decade vancomycin resistant *Enterococcus faecium* (VREfm) have become increasingly prevalent causing infections especially in immunocompromised patients.^1,2^ The endemic occurrence of VREfm and increasing number of outbreaks with only limited remaining therapeutical options led to the WHO classification as a high priority pathogen in 2017.^3,4^

The reliable detection of VRE is essential for the initiation of adequate therapy and infection control measures. However, especially low level resistance mediated by *vanB* in *Enterococcus faecium* is difficult to detect (hereafter referred to as “occult VRE”) as the minimal inhibition concentration (MIC) does not necessarily resemble phenotypic vancomycin resistance.^5^ In July 2018 EUCAST issued a warning against the use of vancomycin gradient tests to detect vancomycin resistance in *E. faecium* and *E. faecalis*.^6^

In this prospective study we analyzed if *vanB* mediated vancomycin resistance can be reliably detected by Vitek2 and whether vancomycin resistance can emerge after vancomycin exposure *in vitro*. This is essential for patient management as vancomycin remains the first line antibiotic for treatment of *E. faecium* infections.^7^

## Materials/methods

From 10/2018 until 05/2019 (8 months) 1344 non-copy enterococcal isolates from routine clinical specimens were tested prospectively using the AST-P592 card in a Vitek2 system (bioMérieux, Nürtingen, Germany). Additionally, a bacterial suspension equivalent to 0.5 McFarland standard was inoculated on a chromID VRE screening agar (bioMérieux) and incubated for 48 hours. If vancomycin susceptibility was determined by Vitek2 (MIC ≤ 4mg/L) but growth was detected on the VRE screening agar a PCR for *vanA/vanB* was performed using the GeneXpert *vanA/B* test (Cepheid, Frankfurt, Germany). In case of *vanA* or *vanB* detection, isolates were classified as VRE. For isolates which were positive for *vanA* or *vanB* but phenotypically vancomycin susceptible the vancomycin MIC was further assessed by broth microdilution using the MICRONAUT-S MRSA/GP plate (Merlin Diagnostika, Bornheim, Germany) and vancomycin MIC test strips according to the macromethod (Liofilchem, Roseto degli Abruzzi, Italy) as recommended by the manufacturer and previously described.^8^ For occult VRE, results of susceptibility testing and VRE PCR were also verified at the National Reference Centre for Enterococci at the Robert Koch Institute.

Additionally, to analyze if any VRE were missed using regular susceptibility testing or by VRE screening agar and to ensure methodical consistency, all available *E. faecium* (243/491) from routine clinical specimens isolated between March to May 2019 were analyzed for *vanA/vanB* presence using the Anyplex VanR real-time Detection kit (Seegene, Düsseldorf, Germany), irrespective of whether growth on the VRE screening media was observed.

Furthermore, we compared if follow-up isolates of patients with occult VRE developed phenotypic vancomycin resistance.

## Induction of vancomycin resistance

Phenotypically vancomycin susceptible but *vanA/B* positive isolates were grown in brain-heart-infusion broth supplemented with vancomycin in increasing concentrations (1 mg/L, 2 mg/L, 4 mg/l, 10 mg/l and 25 mg/L) for 48h at 36°C in a rotary shaker at 200 rpm. If growth was detected, blood agar plates were inoculated with broth cultures containing the highest vancomycin concentration. Subsequently, the vancomycin MIC of vancomycin exposed isolates was determined by microdilution and growth on chromID VRE was tested (supplementary fig. 1).

## Results

The majority of screened isolates (63.5%; 853/1344) were *E. faecalis* of which none were VRE (0%) (Figure 1). Therefore, *E. faecalis* isolates were not further investigated. In contrast, for *E. faecium* 156 out of 491 isolates were identified as vancomycin resistant (31.8%) (Figure 1).

**Figure 1:**
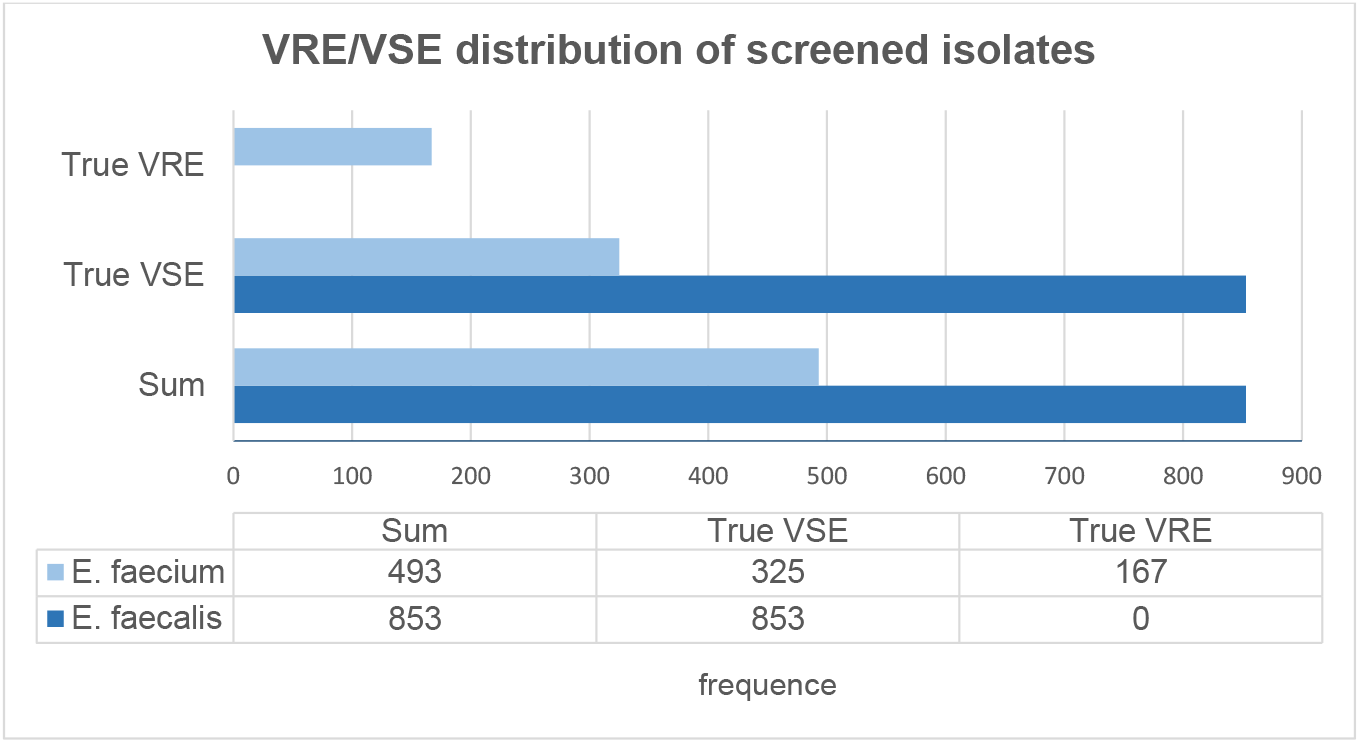
Species distribution of enterococcal isolates in routine clinical specimen (10/2018-05/2019)

Among all tested *E. faecium* isolates, *vanB* was predominant: 87.0% (94/108) were *vanB*, 17.6% (19/108) *vanA* and 2.8% (3/108) possessed both resistance determinants.

Vitek2 did not detect 14 low-level resistant *E. faecium* (occult VRE); of these, three were detected by growth on the VRE screening agar (Table 1). The sensitivity of Vitek2 for VRE detection was 91.0% (142/156) whereas the chromID screening agar’s detection rate was 93.6% (146/156; *p* = 0.52).

**Table 1:** Low-level vancomycin resistance mediated by *vanB* in *Enterococcus faecium* undetected by Vitek2 (occult VRE) before and after vancomycin exposure.

| isolate | before vancomycin exposure |  |  |  |  |  | after vancomycin exposure |  |  |
| --- | --- | --- | --- | --- | --- | --- | --- | --- | --- |
|  | Growth on chromID screening agar | vancomycin MIC by broth microdilution [mg/L] | vancomycin MIC by Vitek2 [mg/L] | vancomycin MIC by MIC test strip [mg/L] |  | Growth in BHI supplemented with vancomycin [mg/L] | Growth on chromID screening agar | vancomycin MIC by broth microdilution [mg/L] | vancomycin MIC by Vitek2 [mg/L] |
|  |  |  |  | 24 h | 48 h |  |  |  |  |
| I-36 | + | ≥32 | **0.5** | 8.0 | 32 | 25 | na | ≥32 | na |
| I-33 | + | ≥32 | **0.5** | 8.0 | 16 | 25 | na | ≥32 | na |
| I-62 | + | 0.5 | **0.5** | 2.0 | 2.0 | 1 | na | 0.5 | na |
| I-81 | - | ≥32 | **1.0** | ≥256 | ≥256 | 25 | + | ≥32 | ≥32 |
| I-78 | - | ≥32 | **0.5** | ≥256 | ≥256 | 25 | + | ≥32 | ≥32 |
| I-43 | - | ≥32 | **0.5** | ≥256 | ≥256 | 25 | + | ≥32 | ≥32 |
| II-61 | - | ≥32 | **0.5** | ≥256 | ≥256 | 25 | + | ≥32 | ≥32 |
| I-45 | - | ≥32 | **0.5** | 32 | 32 | 25 | + | ≥32 | ≥32 |
| I-28 | - | ≥32 | **0.5** | 8 | 32 | 25 | + | ≥32 | 8 |
| I-24 | - | ≥32 | **0.5** | 8 | 16 | 25 | + | ≥32 | ≥32 |
| VIII-44 | - | ≤1 | **0.5** | 2 | 2 | 1 | - | 1 | na |
| I-57 | - | ≤1 | **0.5** | 2 | 2 | 1 | - | 1 | na |
| IV-74 | - | ≤1 | **0.5** | 2 | 2 | 1 | - | 1 | na |
| VI-3 | - | 0.5 | **0.5** | 2 | 2 | 1 | - | 0.5 | na |

na: not applicable

As shown in Table 1, nine of the 14 isolates with low-level vancomycin MICs demonstrated high-level MICs by broth microdilution. The remaining five isolates showed low-level vancomycin MICs both in Vitek2 and broth microdilution (Table 1). Although MICs determined by MIC test strips varied from the MICs by Vitek2 and broth microdilution (Table 1), no very major errors were observed (classification as false vancomycin susceptible).

Prior to an additional MIC screening by broth microdilution, 14 occult VRE were exposed to vancomycin in increasing concentrations. Interestingly, the MIC of exposed strains remained nearly identical to the MIC of the original strains when tested by broth microdilution (Table 1). Seven of 11 occult VRE which initially did not grow on the VRE screening media demonstrated growth after vancomycin exposure. These seven isolates were also showed high-level vancomycin MICs in the broth microdilution and grew in the brain-heart-infusion broth with the maximum vancomycin supplement (25 mg/L).

For eight of the 14 patients with initially occult VRE follow up specimens were available. In four out of these eight patients VRE with high-level vancomycin MICs were detected within one month. Unfortunately, clinical data on antimicrobial therapy of the patients (e.g. glycopeptide therapy) was not available.

## Discussion

Over the last decades, vancomycin resistant *E. faecium* have emerged as a new potentially life-threatening pathogen especially in immunocompromised patients.^1,2^ Due to their significance for therapy and infection control, a reliable detection of this pathogen is essential for patient health. In this real-life prospective study, we demonstrate that VREfm frequently remain undetected by Vitek2 and chromID VRE. Additionally, not only low-level resistant VREfm remain undetected but also MICs of high-level vancomycin resistant VREfm (HLVRE) were frequently underestimated (9/14 isolates).^8^ All isolates not identified as VRE by Vitek2 were *vanB* type (14/14).

Interestingly, in isolates possessing *van* genes but without phenotypic vancomycin resistance, MICs did not increase significantly after exposure to vancomycin. We expected that vancomycin exposure may lead to an enhanced induction of the *vanH_B_BX_B_* operon, resulting in development of phenotypic resistance against vancomycin, as previously reported for vancomycin-variable *E. faecium* (VVE).^9–12^ The occult VRE in the present study were unlike VVE which are *vanA* VREfm reverting from a VSE phenotype to VRE by constitutive expression of the vancomycin resistance cassette.^9^ VVE possess *vanA*, demonstrate low vancomycin MICs, do not grow on VRE screening media and may cause treatment failure if treated with vancomycin.^9–11,13^ In 2018, Hashimoto *et al*. described a stealthy *vanB* cluster which consisted of vancomycin susceptible *E. faecium* (MIC = 3 mg/L) that possessed the ability to spontaneously revert to a resistant phenotype mainly caused by a change of *vanB* gene transcription levels.^14^ The majority of our occult VRE only demonstrated low-level MICs by Vitek2 but high MICs by broth microdilution or MIC test strips indicating an existing high-level vancomycin resistance. Unlike stealthy VRE^14^ or vancomycin-variable VRE^9–11^ occult VRE were mainly VRE evading detection by Vitek2.

As the MIC determination by Vitek2 is merely a deduction of three antibiotic concentrations in comparison to reference data instead of a single endpoint reading, the uniqueness of occult VRE may not be based on a variation of the *van* gene but on their growth capacity. Vitek2’s inadequacy to detect occult VRE may therefore be based on inapt reference data. As undetected VRE pose a threat for treatment failure for patients with VRE infections possible evasion mechanisms need to be further investigated.

The clinical significance of the *van* gene detection in the five isolates which were *vanB* positive but had low vancomycin MICs despite repeated exposure to vancomycin remains unclear. Interestingly, we identified 4 cases who initially had occult VRE and within one month developed VRE which were detected by Vitek. Unfortunately, it was not possible to collect and compare these isolates to the original strains.

Until now, to the best of our knowledge, no data is available on the outcome of patients infected by occult VRE who are treated with vancomycin. In an outbreak among neonatal patients in Germany the existence of phenotypic and genotypic discrepancies in *vanB* positive *E. faecium* was reported. The outbreak was only detected fortuitously and a colonization rate between 10-25% was assumed.^15^ In this study only specimen associated with clinical infection were included and a prevalence of almost 10% was recorded. The colonization with occult VRE is bound to be higher and may be the reason for the extensive spread of *vanB* positive VRE in Germany beginning in 2007.^4,15^

In conclusion, our data demonstrates the inability of Vitek2 to reliably detect even high-level vancomycin resistance in *E. faecium* that is associated with the *vanB* genotype. Additional methods (e.g. culture on VRE screening agar or PCR) are necessary to improve detection of *van*-positive enterococci in clinical routine and to avoid treatment failure. As *vanA/B* PCR for all clinical *E. faecium* isolates is neither feasible nor cost-efficient in the setting of a routine lab, subculture on VRE screening media represents a pragmatic solution and increases VRE detection rates. Therefore, further studies are needed to determine which screening agar performs best for VRE detection in combination with Vitek2. Considering that 9% of the VRE in this study were occult VRE we recommend performing *vanA/B* PCR of all *E. faecium* isolated from invasive infections (including all blood cultures).

## Acknowledgements

We thank all technicians of the diagnostic laboratory of the Institute for Medical Microbiology, Immunology and Hygiene and at the National Reference Centre for excellent technical assistance.

## Transparency declaration

The authors have nothing to declare.

## References

1. Arias CA, Murray BE. 2012. The rise of the Enterococcus: beyond vancomycin resistance. Nat. Rev. Microbiol. 10, 266–278

2. Miller WR, Murray BE, Rice LB, Arias, CA. 2016. Vancomycin-Resistant Enterococci: Therapeutic Challenges in the 21st Century. Infect. Dis. Clin. North Am. 30, 415–439

3. Frakking FNJ, Bril WS, Sinnige JC, Klooster JEV, de Jong BAW, van Hannen EJ, Tersmette M. 2018. Recommendations for the successful control of a large outbreak of vancomycin-resistant Enterococcus faecium in a non-endemic hospital setting. J. Hosp. Infect. 100, e216–e225

4. WHO publishes list of bacteria for which new antibiotics are urgently needed. https://www.who.int/news-room/detail/27-02-2017-who-publishes-list-of-bacteria-for-which-new-antibiotics-are-urgently-needed (last accession date: 10/21/2020)

5. Klare I, Fleige C, Geringer U, Witte W, Werner G. 2012. Performance of three chromogenic VRE screening agars, two Etest(®) vancomycin protocols, and different microdilution methods in detecting vanB genotype Enterococcus faecium with varying vancomycin MICs. Diagn. Microbiol. Infect. Dis. 74, 171–176

6. EUCAST Warning, 10.07.2018, http://www.eucast.org/fileadmin/src/media/PDFs/EUCAST_files/Warnings/Warning_against_Low_MIC_VanB_2018.pdf. (last accession date: 04/05/2019)

7. Doernberg SB, Lodise TP, Thaden JT, Munita JM, Cosgrove SE, Arias CA, Boucher HW, Corey GR, Lowy FD, Murray B, Miller LG, Holland TL; Gram-Positive Committee of the Antibacterial Resistance Leadership Group (ARLG). 2017. Gram-Positive Bacterial Infections: Research Priorities, Accomplishments, and Future Directions of the Antibacterial Resistance Leadership Group. Clin Infect Dis. 2017 Mar 15;64(suppl_1):S24–S29.

8. Klare I, Bender JK, Fleige C, Kriebel N, Hamprecht A, Gatermann S, Werner G. Comparison of VITEK® 2, three different gradient strip tests and broth microdilution for detecting vanB-positive Enterococcus faecium isolates with low vancomycin MICs. 2019. J. Antimicrob. Chemother. 74, 2926–2929

9. Thaker MN, Kalan L, Waglechner N, Eshaghi A, Patel SN, Poutanen S, Willey B, Coburn B, McGeer A, Low DE, Wright GD. Vancomycin-variable enterococci can give rise to constitutive resistance during antibiotic therapy.2015. Antimicrob. Agents Chemother. 59, 1405–1410

10. Kohler P, Eshaghi A, Kim HC, Plevneshi A, Green K, Willey BM, McGeer A, Patel SN, for the Toronto Invasive Bacterial Diseases Network (TIBDN). 2018. Prevalence of vancomycin-variable Enterococcus faecium (VVE) among vanA-positive sterile site isolates and patient factors associated with VVE bacteremia. PloS One 13, e0193926

11. Sivertsen A, Pedersen T, Larssen KW, Bergh K, Rønning TG, Radtke A, Hegstad K. 2016. A Silenced vanA Gene Cluster on a Transferable Plasmid Caused an Outbreak of Vancomycin-Variable Enterococci. Antimicrob. Agents Chemother. 60, 4119–4127

12. Hill CM, Krause KM, Lewis SR, Blais J, Benton BM, Mammen M, Humphrey PP, Kinana A, Janc JW. 2010. Specificity of Induction of the vanA and vanB Operons in Vancomycin-Resistant Enterococci by Telavancin. Antimicrob. Agents Chemother. 54, 2814–2818

13. Oravcova V, Kolar M, Literak I. 2019. Highly variable vancomycin-resistant enterococci in the north-eastern part of the Czech Republic. Lett. Appl. Microbiol. 69, 16–22

14. Hashimoto Y, Kurushima J, Nomura T, Tanimoto K, Tamai K, Yanagisawa H, Shirabe K, Ike Y, Tomita H. 2018. Dissemination and genetic analysis of the stealthy vanB gene clusters of Enterococcus faecium clinical isolates in Japan. BMC Microbiol. 18, 213

15. Werner G, Klare I, Fleige C, Geringer U, Witte W, Just HM, Ziegler R. 2012. Vancomycin-resistant vanB-type Enterococcus faecium isolates expressing varying levels of vancomycin resistance and being highly prevalent among neonatal patients in a single ICU. Antimicrob. Resist. Infect. Control 1, 21

